# Measuring urinary cortisol and testosterone levels in male Barbary macaques: A comparison of EIA and LC–MS

**DOI:** 10.1101/506923

**Authors:** Alan V. Rincon, Julia Ostner, Michael Heistermann, Tobias Deschner

**Affiliations:** Department of Behavioral Ecology, Johann-Friedrich-Blumenbach Institute for Zoology and Anthropology, University of Goettingen, Goettingen, Germany; Research Group Social Evolution in Primates, German Primate Center, Leibniz Institute for Primate Research, Goettingen, Germany; Leibniz ScienceCampus Primate Cognition, Goettingen, Germany; Endocrinology Laboratory, German Primate Center, Leibniz-Institute for Primate Research, Goettingen, Germany; Department of Primatology, Max Planck Institute for Evolutionary Anthropology, Leipzig, Germany

**Keywords:** Testosterone, Cortisol, EIA, LC–MS, Urine, Barbary macaque

## Abstract

The development of methods to quantify hormones from non-invasively collected samples such as urine or feces has facilitated endocrinology research on wild-living animals. To ensure that hormone measurements are biologically meaningful, method validations are highly recommended for each new species or sample matrix. Our aim was to validate three commonly used enzyme immunoassays (EIA), one for analysis of cortisol and two for analysis of testosterone, to assess adrenocortical and gonadal activity, respectively, from the urine of male Barbary macaques. We compared EIA and liquid chromatography–mass spectrometry (LC–MS) results to determine if the EIA measurements truly reflect levels of the target hormone and to determine if cross-reactivities with other steroids were potentially confounding results. Furthermore, we conducted a biological validation of testosterone to ensure that both EIA and LC–MS were able to capture physiologically meaningful differences in hormone levels. We found that cortisol measured by EIA correlated strongly with cortisol measured by LC–MS in both adult and immature males, without the need for deconjugation of steroids in the urine. Both testosterone EIAs correlated strongly with LC–MS in adult males, but only if steroids in the urine were deconjugated by enzymatic hydrolysis prior to analysis. However, in immature males, EIA and LC–MS results did not correlate significantly. Further correlation analyses suggest this is likely due to cross-reactivity of the testosterone antibody with other adrenal steroids such as cortisol, DHEA, and likely others, which are present at much higher concentrations relative to testosterone in immature males. Testosterone levels were significantly higher in adult compared to immature males as measured by LC–MS but not as measured by EIA. Taken together, our results suggest that the testosterone EIAs are suitable to assess gonadal activity in adult but not immature males, and only if a hydrolysis of the urine is conducted prior to analysis.

## 1 Introduction

Steroid hormones facilitate a range of behaviors and developmental changes in animals. For example, glucocorticoids help to mobilize energy reserves and respond adaptively to environmental and social stressors (Beehner and Bergman, 2017). Androgens, such as testosterone (T), the predominant male sex-hormone, promote the production of sperm, development of secondary sexual characteristics, and male reproductive competition (Wingfield et al., 1990). To study steroid hormones in wild-living animals, behavioral ecologists and wildlife endocrinologists are increasingly measuring hormone levels non-invasively, usually via the analysis of excreted hormone metabolites in urine or feces (Behringer and Deschner, 2017; Palme, 2019). This is in part due to the practical and ethical advantages these methods have compared to the traditional approach of measuring hormones invasively from blood. Notably, urine and fecal samples can be collected repeatedly over time without the need to capture or disturb the animal.

The most commonly used methods to quantify hormone concentrations in the various matrices are radio-(RIA) and enzyme-(EIA) immunoassays. These methods rely on the use of antibodies that bind to the hormone of interest, thus allowing for its concentration to be quantified in a sample (Grange et al., 2014). Despite their specificity, antibodies used in immunoassays may cross-react with other structurally similar metabolites. In blood, where steroid hormones circulate in their free biologically active form at much higher concentrations than its metabolites, the impact of such cross-reactivities is often negligible. However, steroids are extensively metabolized in the liver and/or by gut bacteria (Palme, 2019), and as a result, the concentration of the free biologically active hormone excreted in urine or feces is usually very low relative to its metabolites or conjugated forms (Bahr et al., 2000; Möhle et al., 2002; Palme and Möstl, 1997). Thus the impact of cross-reactivity is usually more pronounced in urine or fecal samples than in blood samples. Nevertheless, if the cross-reacting metabolites originate from the parent hormone of interest, the signal detected by the assay may still be biologically meaningful. However, if the cross-reacting compounds measured by the antibody used originate from hormones with different biological functions, then results are confounded and may be uninterpretable. For example, some testosterone immunoassays co-measure androgen metabolites of non-gonadal origin (likely from dehydroepiandrosterone (DHEA), which is of adrenal origin) to such an extent, that they fail to find the predicted difference in testosterone levels between males and females (Goymann, 2005; Möhle et al., 2002). Similarly, two out of four glucocorticoid EIAs showed substantial cross-reactivity with testosterone metabolites in the urine and feces of male African elephants (*Loxodonta africana*), potentially leading to a confound when applied for the assessment of adreonocortical activity in this species (Ganswindt et al., 2003).

As steroids are primarily excreted in their conjugated form in urine (Bahr et al., 2000; Möhle et al., 2002), results from hormone-specific assays designed to measure the free hormone in blood may be improved by first deconjugating the steroids via hydrolysis and/or solvolysis (Hauser, Deschner, et al., 2008; Preis et al., 2011), thus increasing the ratio of free hormone in the sample. Failure to do so may produce inconsistent results. For example, two studies on wild chimpanzees (*Pan troglodytes*) tested for a relationship between dominance rank and urinary testosterone levels. One study found a significant positive correlation, where high ranking males had higher testosterone (Muller and Wrangham, 2004), whereas the other study did not find a significant relationship (Sobolewski et al., 2013). While this could be a true difference between populations, methodological differences in hormone analysis could also account for this discrepancy. The key difference is that in the former study, steroids in the urine were deconjugated via hydrolysis prior to analysis (Muller and Wrangham, 2004), but in the latter study, testosterone was analyzed from unprocessed urine samples (Sobolewski et al., 2013). Considering that the vast majority of testosterone is excreted as glucuronide conjugates in chimpanzee urine (Möhle et al., 2002), it is unclear whether using a testosterone EIA on non-hydrolyzed urine produces biologically meaningful results, particularly when cross-reactivity of the antibody with conjugated testosterone is not known or very low.

Given the discrepancies mentioned above, it is of paramount importance to validate hormone assay methods prior to their application in non-invasively collected samples (Heistermann et al., 2006; Möhle et al., 2002; Touma and Palme, 2005). Furthermore, due to variation in metabolism, method validations are recommended for each new species, sex, or sample matrix (Buchanan and Goldsmith, 2004; Goymann, 2005; Heistermann et al., 2006; Palme, 2019; Touma and Palme, 2005).

Traditionally, two methods have been used to validate measurements of immunoassays. First, radioinfusion studies work by injecting a small amount of radio-labelled hormone into the animal and collecting all subsequent excreta. Since the injected hormone is radio-labelled, researchers are able to deduce the time-lag to hormone excretion, the metabolism pathway, and whether an antibody really cross-reacts with the target hormone (Goymann, 2005; Palme, 2019). Second, physiological validations of steroids may be conducted by pharmacologically inducing their release in the body, then checking if the immunoassay is able to capture the resulting change in hormone levels (Goymann, 2005; Palme, 2019). The downside to both of these methods, however, is that they must be conducted invasively and usually in captivity, neither of which may be practical.

Liquid chromatography–mass spectrometry (LC–MS) can also be used as a tool to validate immunoassay measurements (Gesquiere et al., 2014; Habumuremyi et al., 2014; Preis et al., 2011). In contrast to immunoassays, LC–MS allows for highly specific measurements of hormones in samples based on their molecular weight and charge (Cross and Hornshaw, 2016; Hauser, Deschner, et al., 2008) without any confounding effects of antibody cross reactivity. Therefore, comparing immunoassay measurements to LC–MS is a useful way to deduce if measurements from an immunoassay indeed reflect the concentration of the target hormone and which cross-reacting metabolites may be potentially confounding results (Gesquiere et al., 2014; Habumuremyi et al., 2014; Preis et al., 2011). Both immunoassays and LC–MS may be applied to non-invasively collected samples and the comparison of their measurements offer a useful alternative when more invasive validation methods are not feasible or desirable.

One way to ensure that non-invasive methods can capture natural variations in hormone levels is to conduct biological validations. For example, a testosterone assay should be able to differentiate levels of adult males from those of immature males or females (Möhle et al., 2002; Pineda-Galindo et al., 2017), a glucocorticoid assay should be able to detect rises following putatively stressful events such as translocation, capture and restraint (Pineda-Galindo et al., 2017; Touma and Palme, 2005), and estrogen or progesterone assays should be able to detect changes in female reproductive condition (i.e. menstrual cycle, pregnancy: Pineda-Galindo et al., 2017; Fieß et al., 1999).

In this study, we aimed to determine whether three commonly used EIAs, one cortisol and two testosterone, previously used in other nonhuman primates (Bahr et al., 2000; Möhle et al., 2002; Sobolewski et al., 2013) were suitable to assess adrenocortical and gonadal activity, respectively, in the urine of male Barbary macaques (*Macaca sylvanus*). In this species, validations have been conducted for assays measuring glucocorticoid (Heistermann et al., 2006) and androgen (Rincon et al., 2017) metabolite levels in fecal samples. Only one study has measured cortisol levels in the urine of (female) Barbary macaques (Sonnweber et al., 2015), although no validation has yet been conducted for this species. First, we examined the pattern of conjugation of cortisol and testosterone in the urine of adult and immature male Barbary macaques. Then, we compared the LC–MS cortisol and testosterone results to their respective EIA results from unprocessed urine to determine whether deconjugation steps would be necessary prior to using EIA. If these results did not correlate significantly, we then performed a deconjugation step to see if the correlation improved. The deconjugation step performed (hydrolysis or solvolysis) was chosen based on the pattern of conjugation. To determine any potential influence of cross-reactivity and potential co-measurement of steroid metabolites of different origins on our EIA measurements, we correlated measurements from the three EIAs to cortisol, testosterone and DHEA as measured by LC–MS. To complement the methodological testosterone validations, we additionally performed a biological validation of testosterone by comparing levels of adult males to those of immature males. We predicted that testosterone levels would be higher in adult males compared to immature males (Rincon et al., 2017).

## 2 Materials and Methods

### 2.1 Study site and animals

Study subjects belonged to one out of three groups of Barbary macaques living in free-ranging conditions in 14.5 ha. of enclosed forest at Affenberg Salem, Germany (de Turckheim and Merz, 1984). They are provisioned daily with fruits, vegetables, grains and have *ad libitum* access to water and monkey chow. The study group (group C) consisted of 13-14 adult males, 19 adult females, 2 large sub-adult males, 8 immature males, 10 immature females and 1 newborn infant male. All members of the group were individually recognizable by observers.

### 2.2 Sample collection

A total of 62 urine samples were collected between April and October 2016 from a total of 21 males, including 30 samples from 13 adults (7 to 25 years old), and 32 samples from 8 immature individuals (1 to 4.5 years old). When monkeys were seen to urinate, the urine was caught with a plastic bag when possible or collected from leaves, branches, rocks or the ground by using a disposable pipette or salivette (Salivette Cortisol, Sarstedt, Nümbrecht, Germany). The use of salivettes to collect urine has recently been validated and successfully applied to urine samples from free-ranging macaques (Danish et al., 2015; Müller et al., 2017). Urine samples contaminated with feces were not collected. Urine samples collected by pipette were transferred to 2ml cryotubes. Both samples stored in cryotubes and salivettes were kept in a thermos filled with ice while in the field. At the end of the day, urine was recovered from the salivettes by centrifugation using an electric centrifuge and also transferred to 2ml cryotubes. All samples were then stored in a freezer at −20°C. When data collection was complete, samples were transported in containers with dry ice to the endocrinology laboratory where they were once again kept frozen at −20°C until hormone analysis.

### 2.3 Hormone analysis

#### 2.3.1 Deconjugation and extraction of steroids

The deconjugation and extraction of steroids for LC–MS analysis followed a modified version of a protocol previously described (Hauser, Deschner, et al., 2008; Preis et al., 2011). We used 20 μl urine for analysis. We also added 50 μl internal standard mixture, to control for losses during extraction and purification, and matrix effects on ionisation of MS measurements. Internal standard mixtures contained 2 pg/μl each of prednisolon, d4-estrone, d3-testosterone, and d9-progesterone (Dr. Ehrenstorfer, Augsburg, Germany). Steroid glucuronides were hydrolyzed using *β*-Glucuronidase from *Escherichia coli* (activity: 200 U/40μl) (Sigma Chemical Co., St. Louis, MO, USA). Extracts were purified by solid phase extractions (Chromabond HR-X, 30mg, 1ml, Macherey-Nagel, Dueren, Germany) (Hauser, Deschner, et al., 2008). Afterwards, steroid sulfates were cleaved by solvolysis with ethyl acetate/sulphuric acid (200 mg sulphuric acid, 98% in 250 ml ethyl acetate). This solvolysis step was carried out only for the measurements of steroids by LC–MS (and not EIA). Extraction of steroids was carried out with 5 ml tert. butyl methyl ether, evaporated and reconstituted in 100 μl of 30% acetonitrile. Extraction efficiencies for the LC–MS measurements were 81.9% for cortisol, 77.4% for testosterone and 62.7% for DHEA (Hauser, Deschner, et al., 2008).

To determine the pattern of conjugation, we first extracted the urine to obtain the unconjugated fraction, then performed a hydrolysis on the aqueous phase to obtain the glucuronide fraction and finally performed a solvolysis on the remaining aqueous phase to obtain the sulfate fraction. The concentration of each fraction was determined by LC–MS and summed to provide a measure of the total concentration of hormone (cortisol or testosterone). The pattern of conjugation is reported as the percentage of each fraction of the total sum.

To measure testosterone in the urine via EIA, we performed a hydrolysis (but not solvolysis) and extraction as described in section 2.3.1, but without adding the internal standard mixture to the samples. Instead, to assess the efficiency of the combined hydrolysis and extraction procedure we prepared a stock solution of testosterone-glucuronide (Art. No. T-2000; Merck KGaA, Darmstadt, Germany) with a concentration of 50 pg/μl in MeOH/H_2_O (50/50). Aliquots of 0.4 ml each of this stock solution were stored at −20°C until used in the hydrolysis/extraction process as controls. Hydrolysis and extraction of controls were performed as described in section 2.3.1 (see above). Testosterone levels were measured from hydrolyzed controls using T-EIA I as described in section 2.3.3 (see below). To determine the combined hydrolysis/extraction efficiency, testosterone values measured in the controls were divided by the added concentration (50 pg/μl) and expressed as a percentage. The combined hydrolysis/extraction efficiencies ranged from 68% to 82% recovery (N = 11, mean ± SD: 76.6 ± 4.7%).

#### 2.3.2 LC–MS analysis

LC measurements were carried out using a Waters Acquity UPLC separation module equipped with a binary solvent manager and a column oven (Waters, Milford, MA, USA) and separation was performed on a Waters Acquity BEH C18 column (2.1 x 100 mm, 1.7 μm particle diameter). Eluent A was water with 0.1% formic acid and Eluent B was acetonitrile. MS analyses were carried out on a Waters XEVO TQ-S tandem quadrupole mass spectrometer (Micromass, Manchester, UK) with an electro spray interface (ESI) in positive mode (Wessling et al., 2018). The quantitative analysis by LC–MS was realized in the range of 0.03- 100 ng/μl for cortisol and testosterone and 0.5- 100 ng/μl for DHEA (Hauser, Deschner, et al., 2008; Hauser, Schulz, et al., 2008). One sample was excluded due to internal standard loss of >80%. For all other samples (N = 61), internal standard loss was <45%. We examined LC–MS data with MassLynx (Version 4.1; QuanLynx-Software).

#### 2.3.3 EIA

Immunoreactive urinary cortisol (iuC_EIA_) concentrations were determined in unprocessed diluted urine by microtiter plate enzyme immunoassay using an antiserum against cortisol-3-CMO-BSA and biotinylated cortisol as enzyme conjugate (Palme and Möstl, 1997). Prior to analysis, samples were diluted 1:100 to 1:12,800 (depending on sample type and concentration) in assay buffer and duplicate 50μl aliquots of diluted samples and cortisol standard (50 μl, 0.6 - 40 pg/50μl) were combined with labelled cortisol (50 μl) and antiserum (50 μl) and incubated overnight at 4°C. After incubation, the plates were washed four times, 150 μl (~7 ng) of streptavidin-peroxidase (S5512; Sigma-Aldrich Chemie GmbH, Deisenhofen, Germany) in assay buffer was added to each well and the plates incubated at room temperature in the dark for 60 min and then washed again four times. TMB substrate solution (100 μl; 1-Step Ultra TMB, Thermo Fisher Scientific Inc., Rockford, USA) was subsequently added and the plates incubated at room temperature in the dark for another 45 - 60 min. The enzyme reaction was finally stopped by adding 50 μl of 2 M H_2_SO_4_ to each well and absorbance measured at 450 nm (reference 630 nm) in a spectrophotometer (Biotek). Cross-reactivity of the antibody is given in Palme and Möstl (1997). Serial dilutions of samples showed displacement curves which run parallel to the respective standard curve. Assay sensitivity at 90% binding was 0.6 pg. Intra-assay coefficients of variation (CV) of high and low value quality controls were 5.8% (high) and 7.7% (low) while respective figures for inter-assay CVs were 7.4% (high) and 6.2% (low).

For measurement of immunoreactive urinary testosterone (iuT_EIA_) in all sample types (diluted unprocessed urine samples, and samples following extraction, and hydrolysis), we generally applied a testosterone EIA (T-EIA I) using an antiserum that was purchased from Rupert Palme (University of Veterinary Medicine, Vienna, Austria). In addition, a subset of samples was also measured with a second testosterone EIA (T-EIA II) using an antiserum (R156/7) developed by Coralie Munro (Clinical Endocrinology Laboratory, UC Davis, USA). The latter was done in order to examine whether different EIAs designed for the measurement of testosterone would generally provide the same or different results when applied to the measurement of testosterone metabolites in the urine of Barbary macaques. Antibodies were both raised in rabbits against testosterone-3-CMO-BSA (T-EIA I) and testosterone-6-CMO-BSA (T-EIA II). For the assay, in brief, 50 μl aliquots of diluted samples (1:20 to 1:2500, depending on sample type and concentration) and testosterone standard (50 μl, 0.31 - 20 pg/50μl) were combined with HRP-labelled testosterone (50 μl) and antiserum (50 μl) and incubated overnight at 4°C. After incubation, the plates were washed four times, after which TMB substrate solution (100 μl; 1-Step Ultra TMB, Thermo Fisher Scientific Inc., Rockford, USA) was added and the plates incubated at room temperature in the dark for another 45 - 60 min. The enzyme reaction was finally stopped by adding 50 μl of 2 M H_2_SO_4_ to each well and absorbance measured at 450 nm (reference 630 nm) in a spectrophotometer (see above).

Cross-reactivities for T-EIA I were reported in Palme and Möstl (1994), while those for T-EIA II were reported in Kersey et al. (2010). Serial dilutions of samples of both diluted neat urine and urine following hydrolysis and extraction showed displacement curves which run parallel to the respective standard curve in both assays. Assay sensitivities at 90% binding were 0.3 pg for both EIAs. Intra-assay coefficients of variation (CV) of high and low value quality controls were <10% in both assays. Respective inter-assay CVs were 9.0% (high) and 10.7% (low) for T-EIA I and 8.5% (high) and 14.5% (low) for T-EIA II.

Urinary steroid concentrations were corrected for creatinine, measured as described by (Bahr et al., 2000), to account for differences in urine concentration and are expressed as ng/mg creatinine (ng/mg Cr).

In this paper, we refer to the urinary cortisol, testosterone and DHEA measurements by LC–MS as uC_LC–MS_, uT_LC–MS_ and uDHEA_LC–MS_, respectively. By contrast, we refer to immunoreactive urinary cortisol and testosterone measurements by EIA as iuC_EIA_ and iuT_EIA_, respectively. We would like to emphasize the fact that while LC–MS accurately quantifies the levels of each hormone in urine, the measurements from EIA are influenced by cross-reactivity of the antibody with other compounds.

### 2.4 Statistical analysis

All analyses were conducted in R statistical software version 3.5.1 (R Core Team, 2018). For all analyses we used non-parametric tests and computed exact p-values where appropriate (Mundry and Fischer, 1998).

To compare hormone levels measured by EIA and LC–MS and to detect any potential cross-reactivity in the EIAs used we correlated EIA and LC–MS results using a Spearman’s rank correlation. We used a bootstrapping procedure to avoid pseudo-replication due to having multiple samples from each individual. Using an R script, we randomly selected one urine sample from each male and calculated a correlation coefficient across all males. We repeated this procedure 1,000 times, saving all correlation coefficients. If we only had one sample for an individual, then this sample was re-used in all correlations. To avoid the issue of multiple testing we did not calculate p-values. Instead, to infer significance of the correlations, we calculated 95% confidence interval of all correlation coefficients and if the interval did not include 0, we deemed the correlation to be significant. We report the mean correlation coefficient (rho) from all correlations.

To compare testosterone levels between adult (7 to 25 years old) and immature (1 to 4.5 years old) males, we used a Mann-Whitney U test. Mean hormone levels per individual were used when we had multiple samples from the same individual. The significance value was set to p < 0.05.

## 3 Results

The pattern of conjugation of cortisol and testosterone, as revealed by LC–MS, was very similar between adult and immature males. Both steroids were primarily excreted as glucuronides with sulfates making up only a small portion (Fig. 1). For cortisol, a substantial proportion was also excreted unconjugated, i.e., in free form, while free testosterone made up only a small portion of the total amount of urinary testosterone (Fig. 1).

**Fig 1:**
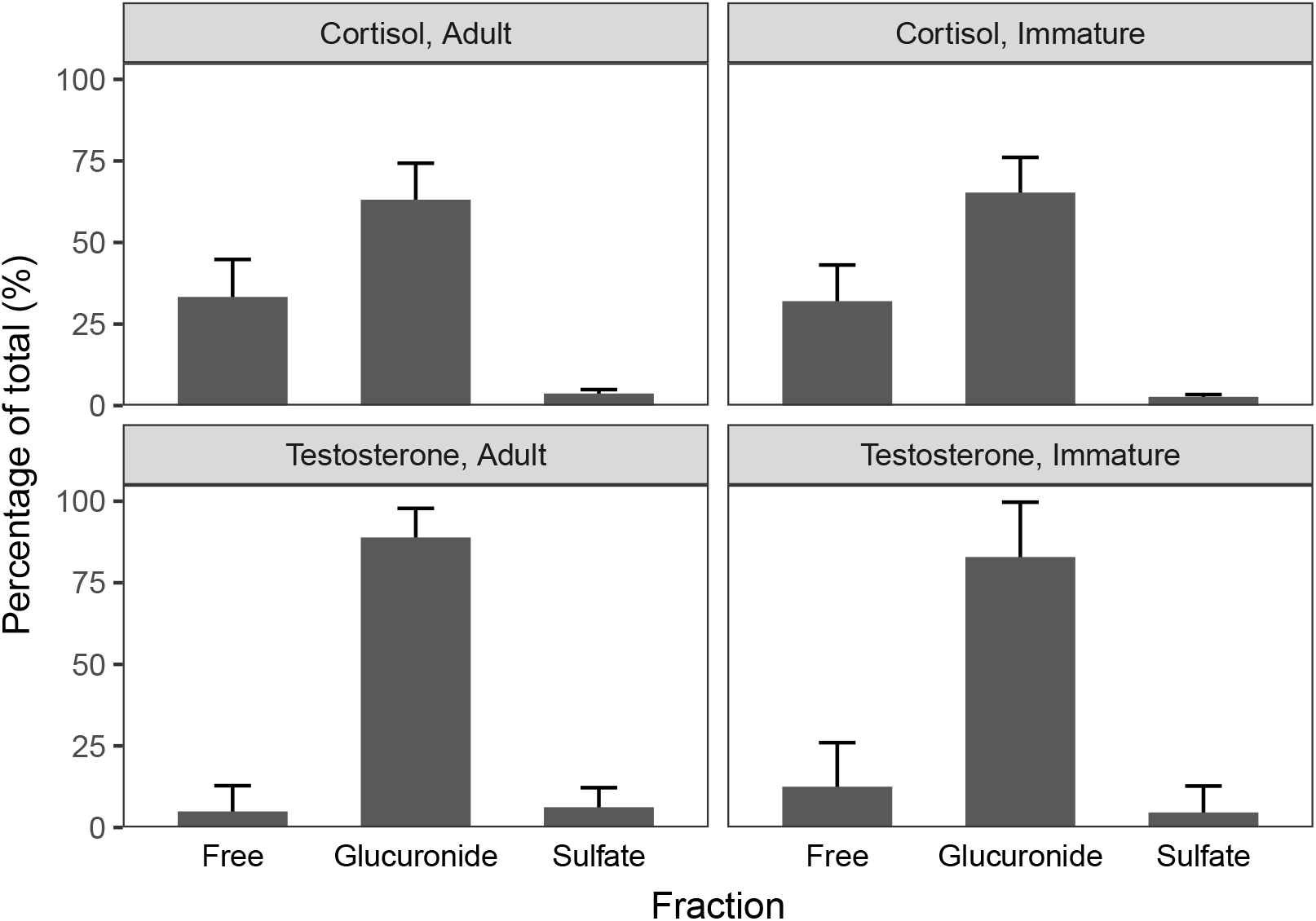
Proportion of cortisol and testosterone excreted in conjugated and unconjugated form in male Barbary macaque urine (as measured by LC–MS). N = 13 adult, N = 8 immature males. Error bars indicate standard deviation.

### 3.1 Steroid levels in urine

For both age classes, uC_LC–MS_ was the most abundant, followed by uDHEA_LC–MS_, and then uT_LC-MS_ (Fig. 2).

**Fig 2:**
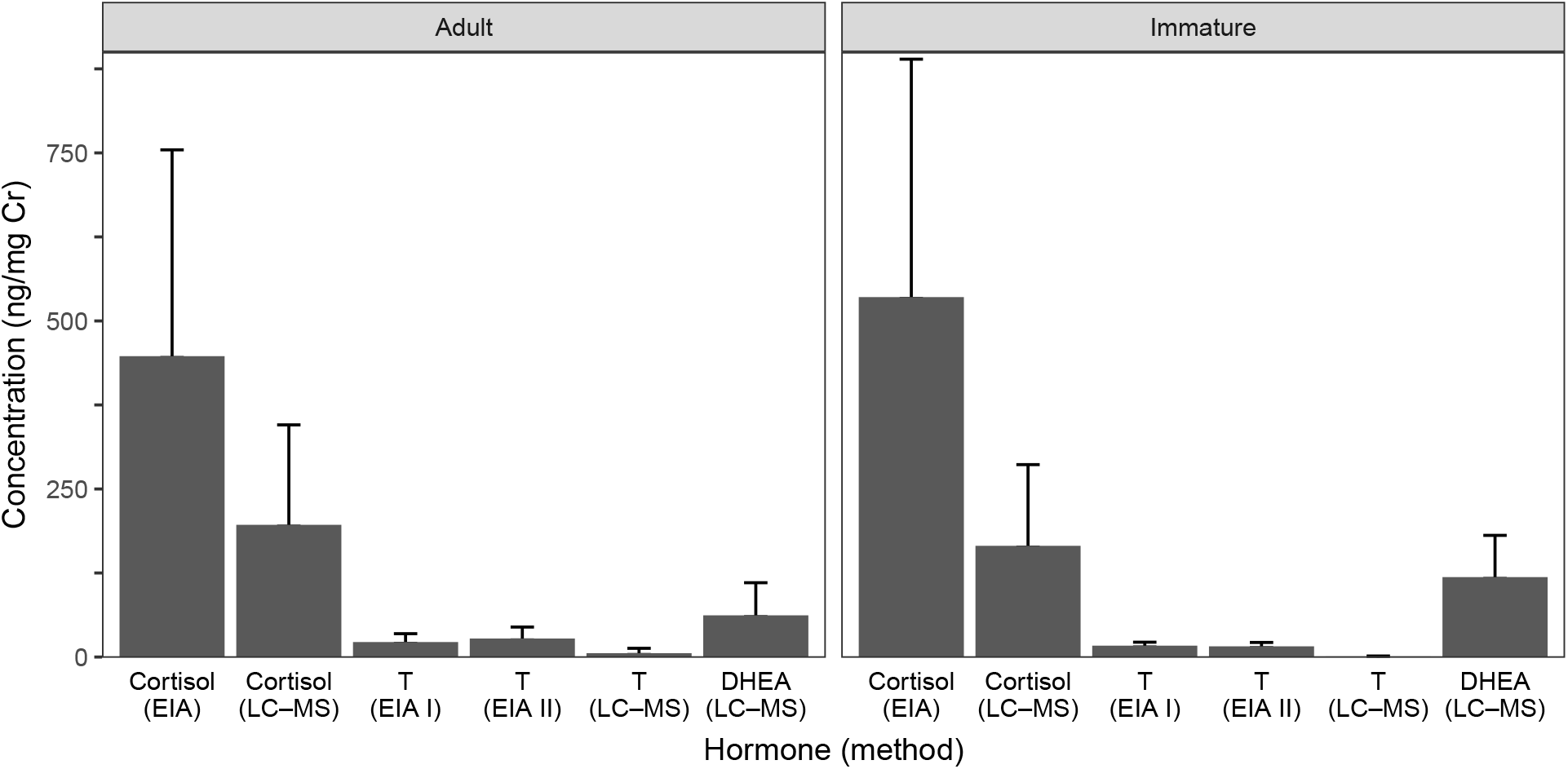
Mean urinary hormone levels as measured by LC–MS and EIA. N = 13 adult (except T-EIA II, N = 10), N = 8 immature males. Error bars indicate standard deviation. Cortisol-EIA used unprocessed urine. Testosterone-EIA used hydrolyzed urine. LC–MS used hydrolyzed and solvolyzed urine.

Absolute concentrations of uC_LC-MS_ and uT_LC-MS_ were higher than iuC_EIA_ and iuT_EIA_, respectively (cortisol adults: 2 times, cortisol immature: 3 times; testosterone adults EIA I: 4 times and EIA II: 5 times, immature EIA I: 19 times, and EIA II: 18 times; Fig. 2). Absolute concentrations of iuT_EIA-I_ and iuT_EIA-II_ were very similar in both adult and immature males (Fig. 2).

### 3.2 Correlations between urinary steroids measured by LC–MS and EIA

iuC_EIA_ as measured from *unprocessed* urine correlated strongly, positively and significantly with uC_LC–MS_ in both adult and immature males (Table 1 and Table 2). This suggests that iuC_EIA_ measurements accurately reflect urinary cortisol levels.

**Table 1:** Bootstrapped Spearman rank correlations (rho) between steroids measured by EIA and LC–MS in adult males. Significant correlations (95% Confidence Interval (CI) does not include 0) in bold.

| EIA | LC-MS | N | $\rho$ | 95% CI | |
| --- | --- | --- | --- | --- | --- |
|  |  |  |  | Lower | Upper |
| Cortisol (unprocessed) | **Cortisol** | **13** | **0.95** | **0.90** | **0.99** |
|  | Testosterone | 13 | -0.02 | -0.35 | 0.30 |
|  | **DHEA** | **13** | **0.54** | **0.31** | **0.73** |
| Testosterone (unprocessed, EIA I) | **Cortisol** | **13** | **0.69** | **0.49** | **0.83** |
|  | Testosterone | 13 | 0.12 | -0.30 | 0.50 |
|  | **DHEA** | **13** | **0.76** | **0.58** | **0.90** |
| Testosterone (unprocessed, EIA II) | **Cortisol** | **9** | **0.35** | **0.12** | **0.58** |
|  | Testosterone | 9 | -0.09 | -0.57 | 0.40 |
|  | **DHEA** | **9** | **0.70** | **0.57** | **0.87** |
| Testosterone (hydrolyzed, EIA I) | Cortisol | 13 | 0.25 | -0.18 | 0.66 |
|  | **Testosterone** | **13** | **0.78** | **0.51** | **0.95** |
|  | DHEA | 13 | 0.23 | -0.10 | 0.52 |
| Testosterone (hydrolyzed, EIA II) | Cortisol | 10 | -0.03 | -0.37 | 0.47 |
|  | **Testosterone** | **10** | **0.76** | **0.45** | **0.95** |
|  | DHEA | 10 | 0.37 | -0.04 | 0.64 |

**Table 2.** Bootstrapped Spearman rank correlations (rho) between steroids measured by EIA and LC–MS in immature males. Significant correlations (95% Confidence Interval (CI) does not include 0) in bold.

| EIA | LC-MS | N | $\rho$ | 95% CI | |
| --- | --- | --- | --- | --- | --- |
|  |  |  |  | Lower | Upper |
| Cortisol (unprocessed) | **Cortisol** | **8** | **0.88** | **0.71** | **1.00** |
|  | Testosterone | 8 | -0.03 | -0.52 | 0.36 |
|  | **DHEA** | **8** | **0.90** | **0.64** | **1.00** |
| Testosterone (unprocessed, EIA I) | **Cortisol** | **8** | **0.73** | **0.43** | **0.93** |
|  | Testosterone | 8 | 0.22 | -0.33 | 0.64 |
|  | **DHEA** | **8** | **0.83** | **0.67** | **0.95** |
| Testosterone (unprocessed, EIA II) | **Cortisol** | **7** | **0.87** | **0.71** | **1.00** |
|  | Testosterone | 7 | -0.12 | -0.68 | 0.39 |
|  | **DHEA** | **7** | **0.85** | **0.64** | **0.96** |
| Testosterone (hydrolyzed, EIA I) | Cortisol | 8 | 0.52 | -0.14 | 0.93 |
|  | Testosterone | 8 | 0.36 | -0.36 | 0.88 |
|  | DHEA | 8 | 0.36 | -0.26 | 0.83 |
| Testosterone (hydrolyzed, EIA II) | **Cortisol** | **8** | **0.83** | **0.62** | **0.98** |
|  | Testosterone | 8 | -0.04 | -0.62 | 0.60 |
|  | **DHEA** | **8** | **0.90** | **0.83** | **0.93** |

iuT_EIA-I_ and iuT_EIA-II_ as measured from *unprocessed* urine did not correlate significantly with uT_LC–MS_ in neither adult nor immature males (Table 1 and Table 2). This suggests that iuT_EIA_ measurements from *unprocessed* urine do not accurately reflect urinary testosterone levels. iuT_EIA-I_ and iuT_EIA-II_ as measured from *hydrolyzed* urine correlated strongly, positively and significantly with uT_LC–MS_ in adult but not in immature males (Table 1 and Table 2). This suggests that iuT_EIA_ measurements from hydrolyzed urine does accurately reflect urinary testosterone levels in adult but not in immature males.

To determine if the cross-reactivity of the cortisol-antibody with testosterone or DHEA were influencing results, we correlated iuC_EIA_ with uT_LC–MS_ and uDHEA_LC–MS_. iuC_EIA_ did not correlate significantly with uT_LC–MS_, in neither adult nor immature males (Table 1 and Table 2). This suggests that cross reaction of the cortisol antibody with urinary testosterone did not have a strong impact on results. However, iuC_EIA_ did correlate significantly with uDHEA_LC–MS_ in both adult and immature males. To tease apart whether this is a cross-reactivity issue or a true correlation between both adrenal steroids, we also correlated uC_LC–MS_ with uDHEA_LC–MS_ and again found a significant positive correlation in both adult and immature males (Bootstrapped Spearman rank correlations: adult: N = 13, rho = 0.40, 95% CI = [0.10, 0.68]; immature: N = 8, rho = 0.70, 95% CI = [0.24, 0.68]). Taken together, this suggests that the cross-reactivity of the cortisol-antibody with DHEA does not significantly impact iuC_EIA_ measurements, but rather that both cortisol and DHEA reflect adrenocortical activity in a similar manner.

To determine if the cross-reactivity of the T-antibodies of EIA I and EIA II with cortisol or DHEA were influencing results, we correlated iuT_EIA-I_ and iuT_EIA-II_ with uC_LC–MS_ and uDHEA_LC–MS_. Both iuT_EIA-I_ and iuT_EIA-II_ as measured from *unprocessed* urine correlated significantly and positively with uC_LC–MS_ and uDHEA_LC–MS_ in both adult and immature males. This suggests that cross-reactivity of both T-antibodies with urinary cortisol and DHEA confound iuT_EIA_ measurements from *unprocessed* urine. iuT_EIA-I_ as measured from *hydrolyzed* urine did not correlate significantly with uC_LC–MS_ or uDHEA_LC–MS_ in neither adult nor immature males. This suggests that the cross-reactivity of the T-antibody of EIA I with urinary cortisol or DHEA does not significantly influence iuT_EIA-I_ measurements in neither adult nor immature males. iuT_EIA-II_ as measured from *hydrolyzed* urine also did not correlate significantly with uC_LC–MS_ or uDHEA_LC–MS_ in adult males. However, it correlated significantly and positively with uC_LC–MS_ and uDHEA_LC–MS_ in immature males. This suggests that the cross-reactivity of the T-antibody of EIA II with urinary cortisol or DHEA does not significantly influence iuT_EIA-II_ measurements in adult males but does significantly influence iuTEIA-II measurements in immature males (Table 1 and Table 2).

### 3.3 Comparison of testosterone levels between age classes

uT_LC–MS_ levels were significantly higher in adult males than immature males (Mann-Whitney U test: N adults = 13, N immature = 8, U = 6, p < 0.001). However, this was not the case for iuT_EIA_ measured from *hydrolyzed* samples (T-EIA I: N adults = 13, N immature = 8, U = 34, p = 0.210; T-EIA II: N adults = 10, N immature = 8, U = 19, p = 0.068).

## 4 Discussion

In this study, we validated the use of cortisol and testosterone EIAs in the urine of adult and immature male Barbary macaques by comparing results to those from LC–MS measurements. iuC_EIA_ measurements from unprocessed urine correlated strongly with uC_LC–MS_ in both adult and immature males. This indicates that the direct measurement of cortisol by EIA is a reliable method for assessing adrenocortical activity in adult and immature male Barbary macaques. iuT_EIA_ measurements only correlated with uT_LC–MS_ if steroids in the urine were enzymatically hydrolyzed prior to analysis in adult males. In immature males, even iuT_EI_A measurements from hydrolyzed urine did not correlate significantly with uT_LC–MS_. This lack of correlation is likely due to cross-reactivity of the testosterone antibody with other adrenal steroids such as cortisol and DHEA, which are present at much higher concentrations relative to testosterone in immature males. Furthermore, uT_LC–MS_ - but not iuT_EIA_ - levels were significantly higher in adult than immature males, biologically validating the measurement of testosterone by LC–MS but not by EIA. Taken together, our results suggest that the testosterone EIAs are suitable to assess gonadal activity from urine analysis in adult but not immature male Barbary macaques, and only if a hydrolysis is conducted prior to analysis.

### 4.1 Pattern of conjugation of cortisol and testosterone

Both cortisol and testosterone were primarily excreted as glucuronide conjugates and only a small proportion was excreted as sulfates. Cortisol was also excreted in substantial quantities in its free form, but testosterone was not. This is consistent with other studies that found that these steroids are primarily excreted in conjugated form in urine in primate (Bahr et al., 2000; Möhle et al., 2002) and non-primate species (Teskey-Gerstl et al., 2000).

### 4.2 Steroid levels in urine

We found that both iuC_EIA_ and iuT_EIA_ levels were significantly higher than uC_LC–MS_ and uT_LC–MS_, respectively, a result consistent with findings from other studies that have compared levels of these two steroids in males from immunoassays and LC–MS (Baid et al., 2007; Hsing et al., 2007; Preis et al., 2011; Welker et al., 2016). Indeed, these “inflation” in absolute hormone concentration as measured by EIA is not surprising given that, in contrast to LC–MS measurements, immunological hormone measurements are generally non-specific as they are influenced by cross-reactivity characteristics of the antibody used. In line with this argument, the difference between EIA and LC–MS measurements is larger in urine samples than in blood samples (Preis et al., 2011). As previously discussed, in blood, steroids primarily circulate in their free biologically active form at much higher proportions than in urine so impact of cross-reactivity of the antibody is lower in blood samples than in urine (Preis et al., 2011). It is important to note that the cross-reactivity of the antibody in immunoassays are not necessarily disadvantageous. As long as the cross-reacting metabolites originate from the same parent hormone, this may actually be advantageous as such so-called group-specific assays may have a higher sensitivity to detect a biologically meaningful change in hormone levels (e.g. Heistermann et al., 2006; Shutt et al., 2012).

### 4.3 Correlations between urinary cortisol measured by EIA and urinary steroids measured by LC–MS

We compared cortisol EIA and LC–MS results to determine if the EIA measurements truly reflect urinary cortisol levels and to determine if cross-reactivities with testosterone or DHEA were confounding results. iuC_EIA_ measured from unprocessed urine correlated strongly and significantly with uc_LC–MS_ for both adult and immature males. This strong correlation suggests that the iuC_EIA_ measurements accurately reflect urinary cortisol levels and was achieved without the need to perform deconjugation steps in the urine samples prior to analysis by EIA. It is likely that cortisol was excreted in sufficiently high quantities in its free form to be measured by the EIA, despite the fact the most of it was excreted as glucuronides. Alternatively, but not mutually exclusive, our cortisol antibody may potentially also cross-react with and measure cortisol in its conjugated forms. The markedly higher absolute levels of cortisol measured by EIA compared to LC–MS (see above) would be in line with such an assumption. Since the deconjugation step is time-consuming and costly, it is worthwhile to check if this analytical step can be avoided prior to any urinary hormone analysis (like in this study) to still produce reliable results.

Cross-reactivities of our cortisol-antibody with urinary testosterone and DHEA were negligible. iuC_EIA_ did not correlate significantly with uT_LC–MS_ in neither adult nor immature males but did have a significant positive correlation with uDHEA_LC–MS_ in both adult and immature males. Rather than this correlation being caused by a cross-reactivity problem, our results indicate that the correlation was caused by simultaneous secretion as uC_LC–MS_ and uDHEA_LC–MS_ were also significantly correlated. The strength of the correlation between uC_LC–MS_ (or also iuC_EIA_) and uDHEA_LC–MS_ was stronger in immature than adult males, possibly indicating a stronger interrelationship between these two adrenal steroids in immature compared to adult males.

Generally, urinary cortisol (or corticosterone) analysis is often used as a measure of adrenocortical activity in various species of mammals (McCallister et al., 2004; McLeod et al., 1996; Palme et al., 1996; Schmid et al., 2001; Touma et al., 2003). In this respect, the cortisol-EIA tested in this study has been shown to successfully pick up the urinary cortisol response to an ACTH challenge in African elephants (Ganswindt et al., 2003). It has also been demonstrated to measure urinary cortisol levels in other nonhuman primates (Bahr et al., 2000), indicating its suitability to non-invasively assess cortisol secretion in various mammalian species. Therefore, we envisage that the measure of urinary cortisol in unprocessed urine as described here is likely suitable for tracking adrenocortical activity in male Barbary macaques, although, to support these results, an accompanying physiological or biological validation would be desirable.

### 4.4 Correlations between urinary testosterone measured by EIA and urinary steroids measured by LC–MS

We compared two testosterone EIAs with LC–MS results to determine if the EIA measurements truly reflect urinary testosterone levels and to determine if cross-reactivity of the antibodies with steroids of adrenal origin, such as cortisol or DHEA, were potentially confounding results. iuT_EIA-I_ and iuT_EIA-II_ measurements from unprocessed urine did not correlate significantly with uT_LC–MS_ in neither adult nor immature males. Since >80% of testosterone was excreted as glucuronide conjugate, we assessed whether enzymatic hydrolysis would result in stronger correlations between our T-EIA and LC–MS measurements. Indeed, this extra analytical step largely improved the correlations between the two data sets so that both iuT_EIA-I_ and iuT_EIA-II_ measurements from hydrolyzed urine now correlated strongly with uT_LC–MS_ in adult males but not immature males. It is likely that cross-reactivity of the testosterone antibodies with cortisol and DHEA, and probably also other adrenal steroids, are confounding the results generated from unprocessed urine in adult and immature males and from hydrolyzed urine in immature males. This is supported by the fact that uC_LC–MS_ and uDHEA_LC–MS_ were both positively correlated with iuT_EIA-I_ and iuT_EIA-II_ when measured from unprocessed urine in both age classes, and with iuT_EIA-II_ - but not iuT_EIA-I_ - when measured from hydrolyzed urine in immature males.

Obviously, the cross-reactivities of the antibodies in the T-EIAs with other metabolites are constant and do not change whether being used with unprocessed or hydrolyzed urine. The observed changes in the strength of correlations with LC–MS measurements are caused by the differences in the ratio of free testosterone to cross-reacting adrenal steroids. For instance, adult males have uC_LC–MS_ and uDHEA_LC–MS_ concentrations approximately 35 and 11 times higher than uT_LC–MS_, respectively (Fig. 2). By contrast, immature males have approximately 186 and 134 times higher uC_LC–MS_ and uDHEA_LC–MS_ levels than uT_LC–MS_, respectively (Fig. 2). In both T-EIAs, the cross-reactivity of the antibody with cortisol and DHEA is <0.1% (Kersey et al., 2010; Palme and Möstl, 1994). Thus, <4% and <1% of the EIA results from hydrolyzed urine can be attributed to cross-reactivity with cortisol and DHEA, respectively, in adult males. This relatively low level of co-measurement of the two adrenal hormones does not obviously confound the T-EIA measurement in adult males as evidenced by the strong positive correlations that both iuT_EIA-I_ and iuT_EIA-II_ have with uT_LC–MS_ when measured from hydrolyzed urine samples. However, in immature males the picture looks different. Around 19% and 13% of the T-EIA results from hydrolyzed urine can be attributed to cross-reactivity with cortisol and DHEA, respectively, a proportion sufficiently large enough to confound testosterone levels measured by EIA in hydrolyzed urine of immature males. Since the vast majority of testosterone was excreted as testosterone glucuronide in male Barbary macaque urine (Fig. 1), the absolute concentration of free testosterone in unprocessed urine was apparently low enough that even in adult males, cross-reactivity with other compounds became a problem. Similar to our results, testosterone immunoassays that have been validated for use in males did not produce reliable results in females, likely due to lower concentration of testosterone and higher proportion of cross-reacting adrenal steroids in females (Goymann, 2005; Preis et al., 2011).

Importantly, cortisol and DHEA are only two examples of hormones coming from a non-gonadal, i.e. the adrenal pathway, whose cross-reaction with the testosterone antibody, potentially affects the measurements of urinary testosterone with the tested T-EIAs. However, there are many other candidate urinary steroid metabolites with a similar potential. For example, one study in chimpanzees found that just among the limited number of potential glucocorticoid and androgen metabolites measured, there were 10 steroids present in the urine of males at higher concentrations than testosterone (Hauser, Deschner, et al., 2008). This potentially leads to a situation where the overall sum of cross-reactivities with all these metabolites, even if the cross-reactivity with each one is very small, will have a much bigger effect than what we describe in this study. This danger is more and more prominent the lower the ratio of the target hormone to confounding metabolites. Concordantly, cortisol, as the most abundant steroid measured in this study, is the least affected by this problem and produced the strongest correlations between EIA and LC-MS measurements. As we show in this study, one approach to mitigate the impact of antibody cross-reactivity in urine samples is to increase the ratio of the target hormone through deconjugation.

### 4.5 Comparison of testosterone levels between age classes

To assess whether our EIA and LC–MS measurements were able to capture biologically meaningful differences in hormone levels, we compared testosterone and DHEA levels of adult and immature males. As predicted, uT_LC–MS_ was significantly higher in adult males compared to immature males, biologically validating the measurement of testosterone by LC–MS in male Barbary macaques. However, there was no significant difference in iuT_EIA_ levels between age classes as measured by either EIAs from hydrolyzed urine. As discussed above, this is likely because in immature males, a substantial co-measurement of adrenal steroids in both T-EIAs increases the concentrations of immunoreactive testosterone and thereby obscures the existing difference in “real” testosterone levels between the two age classes. Similarly to our results, others have found that a testosterone antibody cross-reacting with DHEA metabolites has masked the predicted difference in testosterone levels between adult males and females or castrated males (Goymann, 2005; Möhle et al., 2002). Taken together, our results suggest that measuring testosterone with either EIA from hydrolyzed urine is suitable to assess gonadal endocrine activity in adult but not immature male Barbary macaques. This study highlights that even when a biological validation of a testosterone EIA fails, the tested EIA might still be suitable for specific age and sex classes. At the same time, just because an EIA reliably measures testosterone in one age sex class, e.g. adult males, does not necessarily make it suitable for all other age sex classes. By not only comparing EIA measurements across age sex classes but also correlating EIA with LC–MS measurements we were able to determine in which age class results were reliable and in which they were unreliable.

## 5 Conclusions

Combining LC–MS and EIA approaches, we identified two main issues to consider when measuring steroid hormones from urine samples using immunoassays, i.e. pattern of conjugation and cross-reactivities of the antibody. Both the pattern of conjugation and cross-reactivity are most likely to confound results when the absolute concentration of the target hormone is low relative to other cross-reacting metabolites. As cortisol was excreted in sufficiently high concentrations in its free form in our study subjects, results from EIA and LC–MS were strongly correlated in both adult and immature males, making deconjugation steps unnecessary. Deconjugation of urinary steroid hormones can also be avoided if the antibodies used have a substantial cross-reactivity with the conjugated forms. Testosterone was excreted in urine in much lower concentrations than cortisol and a higher proportion of it was conjugated, primarily as glucuronides, necessitating hydrolysis of steroids to ensure that a strong correlation of EIA compared to LC–MS results was obtained in adult male samples. However, even after hydrolysis, EIA measurements of testosterone in immature male urine samples were still not reliable, most likely due to cross-reactivity with adrenal steroids, which occur at much higher ratio than testosterone in immature males. Taken together, both testosterone EIAs can be used in studies assessing gonadal activity in adult but not immature male Barbary macaques. Even within the same sex, age-related changes in androgen production can cause the same immunoassay to be reliable in one but not another age class. Given differences between steroids in terms of conjugation patterns and absolute levels, preparatory steps in terms of deconjugation and extraction will vary between different EIAs and hormones.

## Acknowledgements

We thank Ellen Merz and Roland Hilgartner for permission to conduct the study. We thank Lauren Cassidy, Tatjana Kaufmann and Lilah Sciaky for help with urine sample collection. We are grateful to Andrea Heistermann and Miriam Polten for support with the EIA measurements and general assistance in the lab, and to Roisin Murtagh and Vera Schmeling for help with the LC–MS analysis. The study benefitted from discussions within the DFG-research group “Sociality and health in primates” (FOR 2136). This research was funded by the Deutsche Forschungsgemeinschaft (DFG, German Research Foundation) - Project number 254142454 / GRK 2070. Consumables for the LC–MS analysis were funded by the Max Planck Society while all reagents for the EIA analyses were funded by the German Primate Center.

